# Real time monitoring of hydrogenotrophic methanogenesis under deep saline aquifers conditions

**DOI:** 10.1101/2025.05.05.652176

**Authors:** Emeline Vidal, Anaïs Cario, Mathilda Jouvin, Maïder Abadie, Olivier Nguyen, Arnaud Erriguible, Anthony Ranchou-Peyruse, Samuel Marre

## Abstract

To investigate the microbial response to deep underground gas injection, specifically CO_2_ and H_2_, a new optically transparent high-pressure reactor was developed to monitor autotrophic microbial growth via *in situ* and *ex situ* characterization techniques. The main advantages rely on avoiding any decompression phases during the entire process, thanks to direct optical access. Here, we monitored the growth of the model methanogenic strain *Methanothermococcus thermolithotrophicus* by applying different H_2_/CO_2_ partial pressures at a total pressure of 100 bar, which is representative of the deep underground storage environment. Additionally, we measured the methane production of the strain at the end of the incubation, which resulted in an increase in methane production with increasing CO_2_ and H_2_ partial pressures until a certain point. These reactors can be used to investigate deep microbial strains under pressure conditions close to their natural environments, eliminating decompression biases.

## Introduction

Since the early twentieth century, the industry has utilized deep geological underground formations for gas storage in various forms, including salt caverns, depleted hydrocarbon reservoirs, and deep aquifers, often alongside indigenous microbial communities ^1,2^. Among these options, porous reservoirs (depleted reservoirs and aquifers) provide the highest storage capacity and host a significant diversity of microbial taxa^3^. Historically, these storage facilities were designed primarily for methane and natural gases^4^. As we entered the 21^st^ century, the substantial rise in anthropogenic CO_2_ emissions and the challenges posed by climate change prompted serious consideration of carbon capture and storage (CCS) strategies, which appear to offer a long-term solution for CO_2_ sequestration in deep geological formations (Carbon Geological Storage – CGS)^5^. More recently, geological storage facilities have been recognized as essential for the emerging dihydrogen (H_2_) sector, presenting a robust, massive and secure storage solution for this energy carrier^6–8^.

Operators and models developers require physico-chemical and microbiological data to ensure these potential future storage sites, which are critical to future energy networks. While studies have demonstrated the ability of methanogenic archaea to convert CO_2_ into methane during enhanced oil recovery (EOR) or CO_2_ geological storage, this phenomenon has generally not been perceived as a concern; quite the opposite^9^. In the context of underground H_2_ storage (UHS), the sustainability of such systems is important. Lithoautotrophic prokaryotes capable of utilizing H_2_ and CO_2_ (*i.e.*, sulfate reducers, methanogens and homoacetogens) exist at storage sites, although their concentrations and activities vary widely^10–16^. Depleted hydrocarbon reservoirs seem particularly suitable for developing underground methanation reactors (UMRs)^17–19^, although some do not display lithoautotrophic properties^20^. While some deep aquifers show significant potential for biomethanation, others do not, suggesting site-specific variability that requires further investigation^2,14–16^. In the UMR context, CO_2_ can originate from several sources: (i) the degradation of organic matter in the reservoir; (ii) the dissolution of carbonate minerals or emissions from deeper geological strata; and (iii) the intentional coinjection of CO_2_ to produce biomethane^9,14,21^.

The future of these storage facilities has generated considerable interest among modelers, yet they work with limited data^22,23^. Experimental data gaps must be filled through multiscale studies from the pore scale^24^ to the pilot scale, high-pressure simulation experiments, and assessments of model microbial strains to establish growth limits and yields, particularly for methanogens^13,25^. Various pathways exist for methanogenesis, including the hydrogenotrophic pathway, which converts H_2_ and CO_2_ into methane^26^.

Research suggests that when H_2_ is injected into deep aquifers, hydrogenotrophic methanogenic populations can develop rapidly, producing substantial amounts of methane. For instance, Amigáň *et al*.^27^ observed 30% consumption of H_2_ within seven months in a deep aquifer used to store town gas (composed of 54% H_2_, 22% CH_4_, and 12% CO_2_ at 40 bar and 25–45°C). Similarly, Haddad and colleagues^14^ simulated 10% H_2_ storage in an aquifer used to store natural gas (99% CH_4_, 1% CO_2_ at 95 bar and 47°C), resulting in a 40% reduction in H_2_ over three months, largely attributed to methanogenesis. Currently, most models rely on the Monod equation, which considers theoretical yields for H_2_ and CO_2_ consumption and methane production^23^. Additionally, the majority of available data in the literature have been gathered at atmospheric pressure and may not accurately reflect the conditions necessary for high-pressure (HP) applications^13,28,29^.

The study of deep microorganisms that grow autotrophically, such as hydrogenotrophic methanogens, poses challenges for HP culture investigations, as these organisms require both aqueous and gaseous (H_2_/CO_2_) phases for growth. Specific cultivation methods have been developed over the last few decades^30–33^, but these methods face several limitations, including (i) the availability of specialized equipment, (ii) the expertise required to utilize this equipment effectively, and (iii) restricted optical access for monitoring growth. Consequently, very few methanogens have been characterized for their piezophilic capabilities^31,33,34^. To cultivate (hyper)thermophilic piezophilic autotrophic methanogens, several strategies have been suggested: (i) use gas-tight syringes with a 2 bar pressure of H_2_/CO_2_ (4:1) in pressurized static pressure vessels^30^; (ii) employ pressure cell reactors with sapphire windows placed in an oven filled with 7.8 bar of H_2_/CO_2_ (4:1) supplemented with helium at the required experimental pressure^31^; and (iii) utilize autoclaves containing cultures in nickel tubes with 2 bar of H_2_/CO_2_ (4:1)^35^. Nevertheless, the decompression step - required for counting cells - can be detrimental and could lead to biases.^36^ Hence, transparent approaches, including high-pressure microfluidics, have been proven to overcome this limitation in monitoring cell growth in real time.^37^

Researchers have generally observed a trend when incubating at very low H_2_/CO_2_ partial pressures while increasing the total pressure of the system, to evaluate the strain tolerance limits to hydrostatic pressure. To assess microbial responses to gas injections effectively in their environment, it is essential to recreate laboratory-scale conditions that mimic those found within the natural ecosystem. The interaction between microorganisms and stored gases is critically important, as it can result in variations in gas quality (including H_2_ consumption and the production of sulfide and/or methane), deterioration of infrastructures (due to biocorrosion), and shifts in physicochemical conditions (affecting water quality and the dissolution or precipitation of biominerals, bioclogging).

In this study, we focus on simulating the physicochemical conditions of a one-kilometer-deep saline aquifer used as either a UHS or a UMR within high-pressure transparent reactors (HPTRs)^5,38–40^. The impact of increasing H_2_/CO_2_ pressure on hydrogenotrophic methanogens is a relatively unexplored area of study. However, given the importance of geological gas storage and the prevalence of these components in the environment, investigating this issue is warranted. One of the novelties of our approach lies in the creation of transparent sapphire cultivation cells that feature a fiber optic system to enable real-time monitoring of microbial growth under conditions simulating various gas injection scenarios. Specifically, this study aims to achieve two objectives: (i) to assess how increases in pressure affect both the growth and metabolism of autotrophic methanogens and (ii) to evaluate the feasibility of biomethane production under conditions that mimic gas injection scenarios by monitoring the kinetics of methane production.

## 1. Materials and methods

### 1.1. Strain and Growth Medium

*Methanothermococcus thermolithotrophicus* strain SN-1 (DSMZ 2095) was used as a model lithoautotrophic thermophilic methanogen^35^. This strain is known for its ability to grow under piezophilic conditions. The strain was cultivated in modified artificial ground water^41^ (AGW) containing NaCl (25.84 g.L^-1^), KCl (0.14 g.L^-1^), MgCl_2_ ·6H_2_ O (1.42 g.L^-1^), CaSO_4_·2H_2_O (1.37 g.L^-1^), CaCl_2_·2H_2_O (0.73 g.L^-1^), NH_4_Cl (0.02 g.L^-1^), yeast extract (1 g.L^-1^) and resazurin (1 mg.L^-1^). To prevent potential acidification during high-pressure experiments, the medium was supplemented with 120 mM HEPES buffer. After autoclaving, the medium was further supplemented with 250 µL of 30 mM K_2_HPO_4_ and 1 mL of SL10 Widdel trace elements^42^. The liquid phase was reduced with 1% (v/v) Na S_2_9H_2_O (25 g.L^-1^). The gas phase was replaced with a mixture of H_2_/CO_2_ at a ratio of 4:1 (Messer) by 15 min of sparging at 1.5 bar. Finally, the pH of the medium was adjusted to 6.8.

### 1.2. High-Pressure Reactors

The high-pressure transparent reactors (HPTRs) utilized in this study are composed of transparent sapphire tubes, each 12 cm long with an internal diameter of 8 mm, providing a total volume of 5 mL (Figure 1-a-b)^43^. Each tube is fitted with titanium alloy (Ta6V) plugs, sealed with Viton® O-rings, and secured by metal clamps. For connectivity, the plugs are machined to accommodate standard commercial VALCO connectors for 1/16’’ tubing (standard 6/32 port). These reactors can withstand pressures of up to 500 bar and temperatures of up to 200°C. The chemical and biological inertness, along with the full transparency of these reactors, allows *in situ* optical measurements such as optical density (OD) (Figure 1-c), facilitating the monitoring of microbial growth without the need to subsample the system under pressure. In certain experiments, internal mixing was achieved by placing a stirring magnet within the growth media and positioning the reactor on a magnetic stirrer.

**Figure 1:**
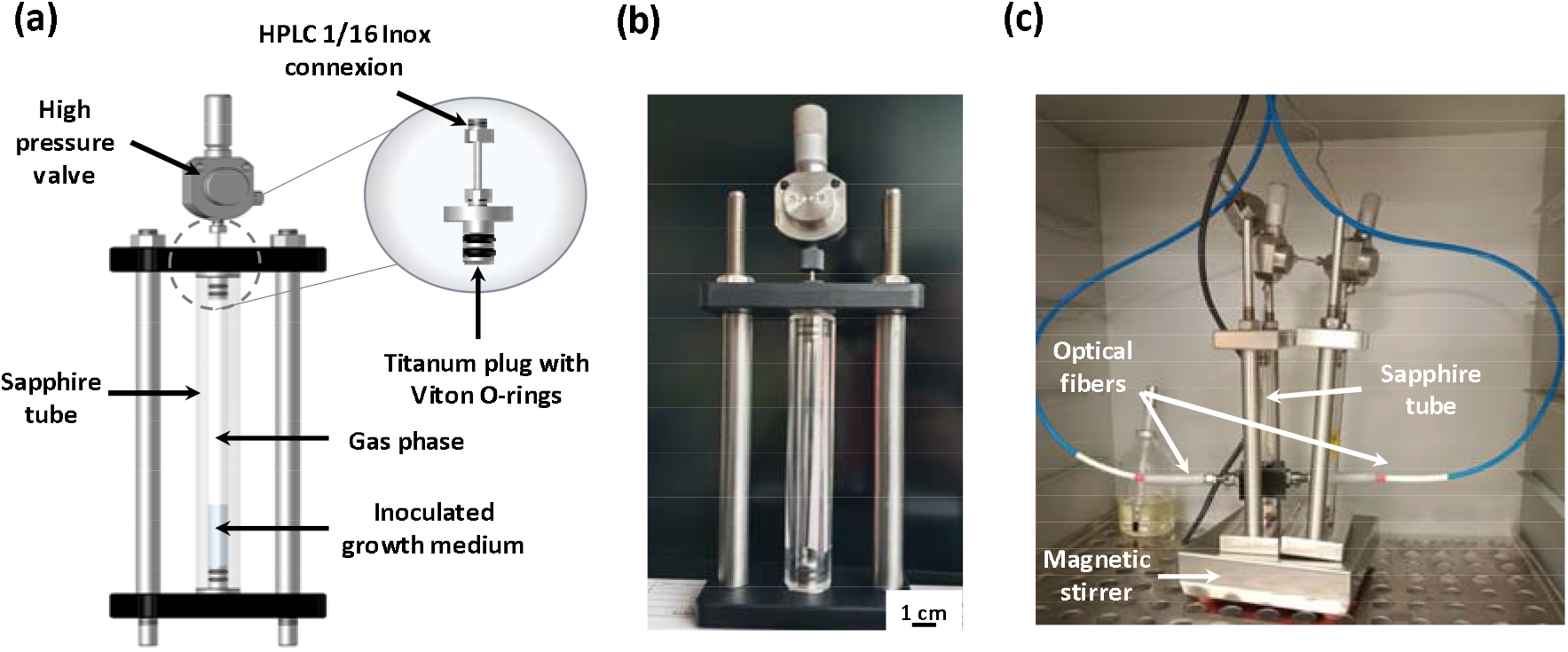
(A) Schematic of the high-pressure transparent reactors (HPTRs) used in this study; (B) photograph of the reactor; and (C) millifluidic sapphire reactor equipped with optical fibers for in situ and real-time O.D. measurements.

### 1.3. High-pressure setup and culture

The archaeal strain was initially cultivated in vials at atmospheric pressure (1.5 bar) and then reinoculated at approximately 2.10^6^ cells.mL^-1^ in fresh medium during the growth phase. Next, 1.5 mL of the inoculated medium was transferred into the HPTRs within an anaerobic chamber (glove box, MB-LABstar, MBraun) under a pure nitrogen atmosphere. The reactors were sealed, removed from the glove box, and immediately incubated at 65°C, the optimum temperature for the strain. The HPTRs were connected to a high-pressure Teledyne ISCO pump (260 HP) filled with a H_2_/CO_2_ mixture (4:1 molar ratio; Messer), adjusted to the desired partial pressure, and supplemented with nitrogen to achieve a total pressure of 100 bar, corresponding to conditions relevant to deep aquifer storage (Figure 2). All the tubing was purged with nitrogen before any injection. Experiments have examined the effects of varying H_2_/CO_2_ partial (p(H_2_/CO_2_)) pressures (5, 15, 20, 30, and 50 bar) on strain growth. Some tests included stirring at 150 rpm via a heat-resistant magnetic stirring device inside an oven (magnetic emotion, MIXdrive 1 eco HT model). All experiments were conducted in triplicate, including a control setup.

**Figure 2.**
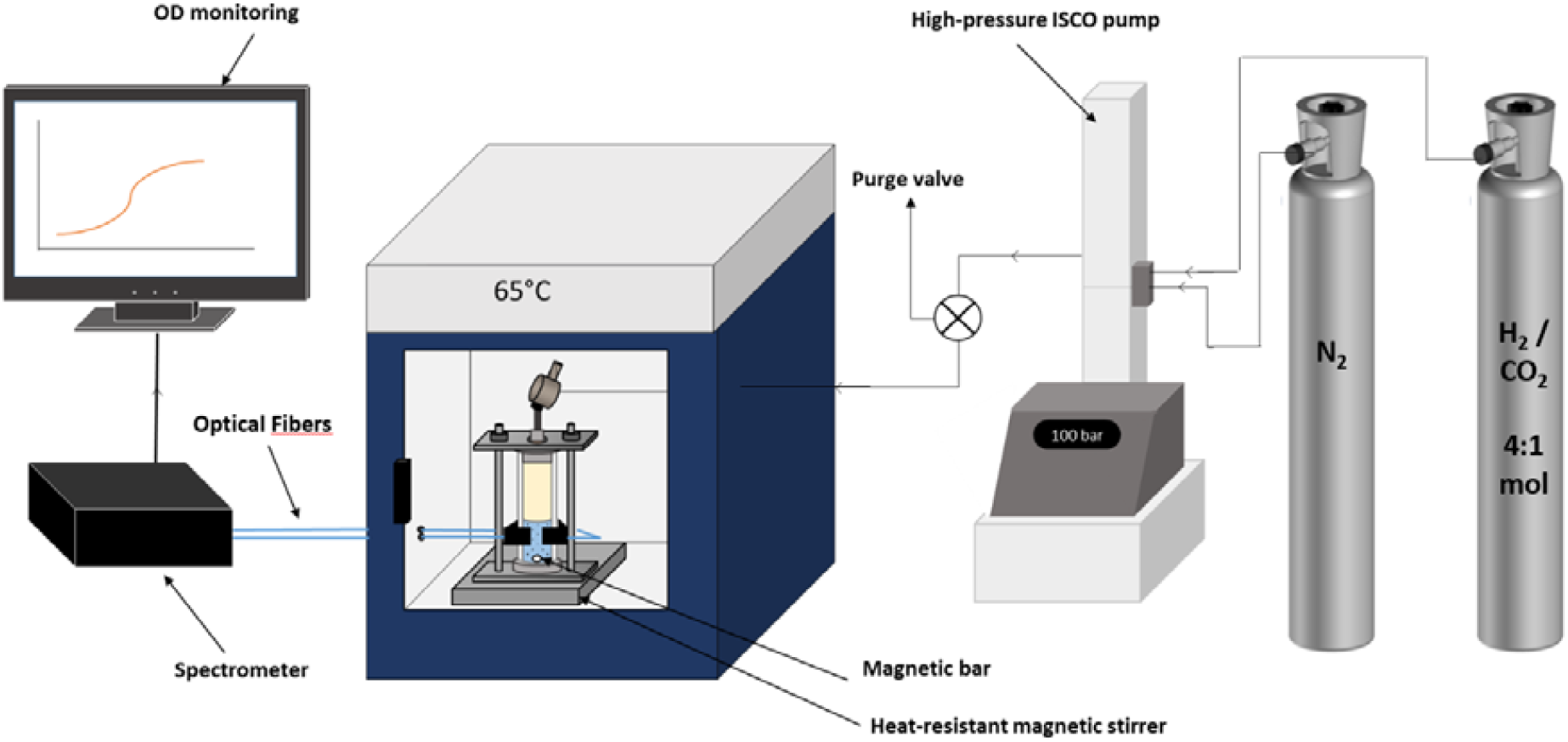
Scheme of the setup used for this study.

### 1.4. Cell Growth Measurements

Optical density (OD) was monitored in real time via optical fibers (IDIL Fibres Optiques, France) connected to the HPTR (Figure 1-c and 2). A spectrometer (Maya 2000 Pro, OceanView software, Ocean Optics) recorded the OD values every second at wavelengths between 600 and 605 nm. Prior to the experiments, validations of the optical system at atmospheric pressure and room temperature confirmed its accuracy; this involved testing with dilutions of stationary phase cultures of *M. thermolithotrophicus*. The initial and final cell concentrations were verified *via* direct cell counting via a Thoma chamber (Preciss, France; surface area: 0.0025 mm^2^; depth: 0.1 mm) under a DM2000 LED phase contrast optical microscope (Leica Microsystems CMS GmbH, Germany). The absorbance of the dilutions was measured at two different wavelengths (*i.e.,* λ = 500 and 600 nm), and the results were similar. The results obtained from the measured values show a linear trend, which validates the system. The validated optic system allowed cell concentrations to be inferred from the recorded optical density (OD) data.

### 1.5. Gas Sampling and Analysis

After the experiment, the methane (CH_4_), dihydrogen (H_2_), carbon dioxide (CO_2_), and nitrogen (N_2_) contents of the gas samples from the HPTRs were analyzed via microgas chromatography (VARIAN CP-4900 PRO Micro-GC) instrument equipped with a thermal conductivity detector and a CP-5A column, with argon used as the carrier gas. A 2 L Tedlar® sampling bag (Supelco) was used to minimize decompression-related artifacts, sampling through a system designed to reduce the pressure drop to 5 bar. A 1/16” 20 cm long tube, with an internal diameter of 500 µm and equipped with two HP valves on both sides, was connected between the HPTRs and the sampling bag. A vacuum-prepared setup enabled sampling of approximately 40 µL of the gas phase at 100 bar, expanding to approximately 4 mL at 1 bar, for analysis. Once recovered, the Tedlar® sampling bag was connected to the micro-GC to be analyzed.

### 1.6. H_2_ and CO_2_ solubility and diffusion modeling

Gaseous substrate solubilities in the liquid phase, including medium salinity effects, were calculated via Crozier and Yamamoto’s model for H_2_^44^ and Duan and Sun’s model for CO_2_^45^, with fugacity computed via their 2006 improvements^46^. For ease of rationalizing the influence of gas penetration in the liquid medium when the system relies only on diffusivity (no stirring, which is not fully accurate since convection occurs in sapphire reactors), we propose analytically estimating the evolution of CO_2_ and H_2_ in the aqueous phase. We assume that the concentration evolves slowly in only one direction (depth of the reactor) under the assumption that convection is negligible. Therefore, the evolution of the concentration is given by the unsteady state mass conservation equation according to the classical Fick’s law for diffusion:

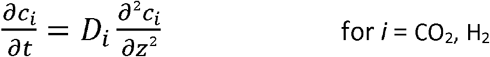

where *c*_*i*_ is the concentration of CO_2_ or H_2_, z is the position and *D*_*i*_ is the diffusion coefficient of CO_2_ or H_2_ in the water phase via the Stokes--Einstein relation at T = 65°C, p_tot_ = 100 bar (*D*_*co2*_= 2.37·10^−9^ m^2^.s^-1^ and *D*_*h2*_ = 8·10^−10^ m^2^.s^-1^). Assuming one-dimensional diffusion in a semi-infinite medium, which is an acceptable hypothesis due to the very low diffusion in water, the well-known analytical solution^47^ is given by:

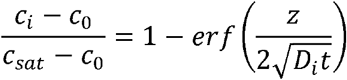

where *c*_*sat*_ and *c*_*0*_ are the initial concentration of CO_2_ or H_2_ in the liquid phase (*c*_*0*_ =0) and the saturation concentration of CO_2_ or H_2_ in the liquid phase, *i.e.*, the concentration values imposed at the boundary conditions z = 0, respectively; *z* is the distance (m); *D*_*i*_ is the diffusion coefficient of molecule *i* in water; and *t* is the time (s). The saturation concentration is given by Henry’s law, which is directly calculated from the partial pressure of the gas in the reactor.

### 1.7. Growth Curves Analysis

The growth curves were analyzed via the R package Growthcurver^48^. Initially, the data were plotted as the average log cell density (LCD). The data were subsequently normalized for each replicate, and the growth rate (R) was subsequently calculated via the following formula:

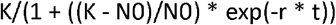

where (N(t)) represents the cell concentration over time, (K) represents the maximum cell population, (N0) represents the initial cell population, (r) represents the growth rate, and (t) represents the time in hours. All raw data are provided in the supplemental material (ESI-1).

## 2. Results and Discussion

Initially, high-pressure transparent reactors (HPTRs) were stirred at 150 rpm to facilitate thermodynamic equilibrium between the gas and liquid phases across a water/liquid interface of 50 mm^2^ and to eliminate concentration gradients. The primary objective of this step was to assess the intrinsic growth potential and production yield of *Methanothermococcus thermolithotrophicus under* varying H_2_ and CO_2_ partial pressures (p(H_2_/CO_2_)). In the second phase, stirring was removed to simulate better environmental conditions, relying solely on diffusivity and slow convective mixing, which is the major transfer mechanism in porous environments.

### 2.1. Effects of p(H_2_/CO_2_) on growth and metabolism

#### 2.1.1. Cell development

In the stirred system, growth kinetics at different partial pressures were monitored via an optical fiber system for *in situ* optical density (OD) measurements. As illustrated in Figure 3, cell growth was observed for p(H_2_/CO_2_) values ranging from 5 to 20 bar, with the highest cell density (1·10^8^ cell.mL^-1^) achieved at 15 bar. No growth was detected at p(H_2_/CO_2_) > 30 bar within the 24-h cultivation period. Comparatively, experiments by Haddad and colleagues^14^ simulated the physicochemical conditions of underground gas storage (UGS) in a deep aquifer with low agitation (20 rpm). The tested H_2_ concentration of approximately 10 bar closely aligned with the optimal condition, from the methanogenesis point of view, established in the current study (p_H2_ = 12 bar). These results warrant further investigation over extended incubation periods (*e.g.*, 5 to 7 days), as higher p(H_2_/CO_2_) levels (30 and 50 bar) may result in longer lag phases for the strain.

**Figure 3.**
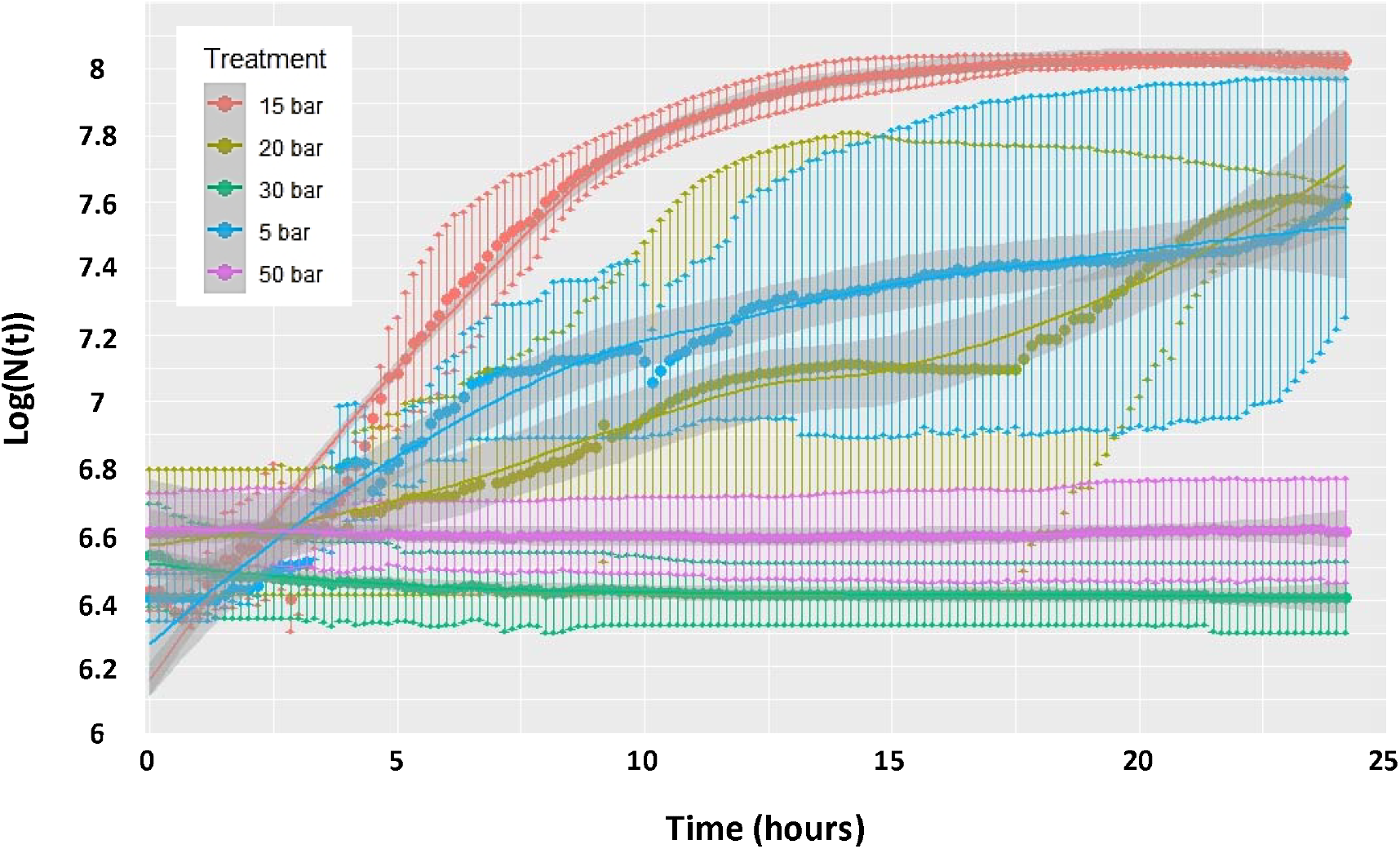
Methanothermococcus thermolithotrophicus growth in response to applied p(H_2_/CO_2_) (80/20 mol%) at a total pressure of 100 bar over 24 hours of incubation. Each curve represents the average of two in situ OD measurements (Log (N(t))).

The specific growth rates for each condition are displayed in Figure 4. Interestingly, the highest growth rate (0.84 ± 0.27 h^-1^) was observed at 20 bar, not at 5 bar (0.57 ± 0.34 h^-1^) or 15 bar (0.60 ± 0.14 h^-1^), despite 15 bar yielding the highest cell density. This suggests that while higher p(H_2_/CO_2_) may accelerate cell growth and production, they do not necessarily correlate with maximum density or methane production. Although an increase in substrates (H_2_ and CO_2_) could imply enhanced growth and production, the optimal cell density at p(H_2_/CO_2_) = 15 bar may indicate potential toxicity from one or both gases.

**Figure 4.**
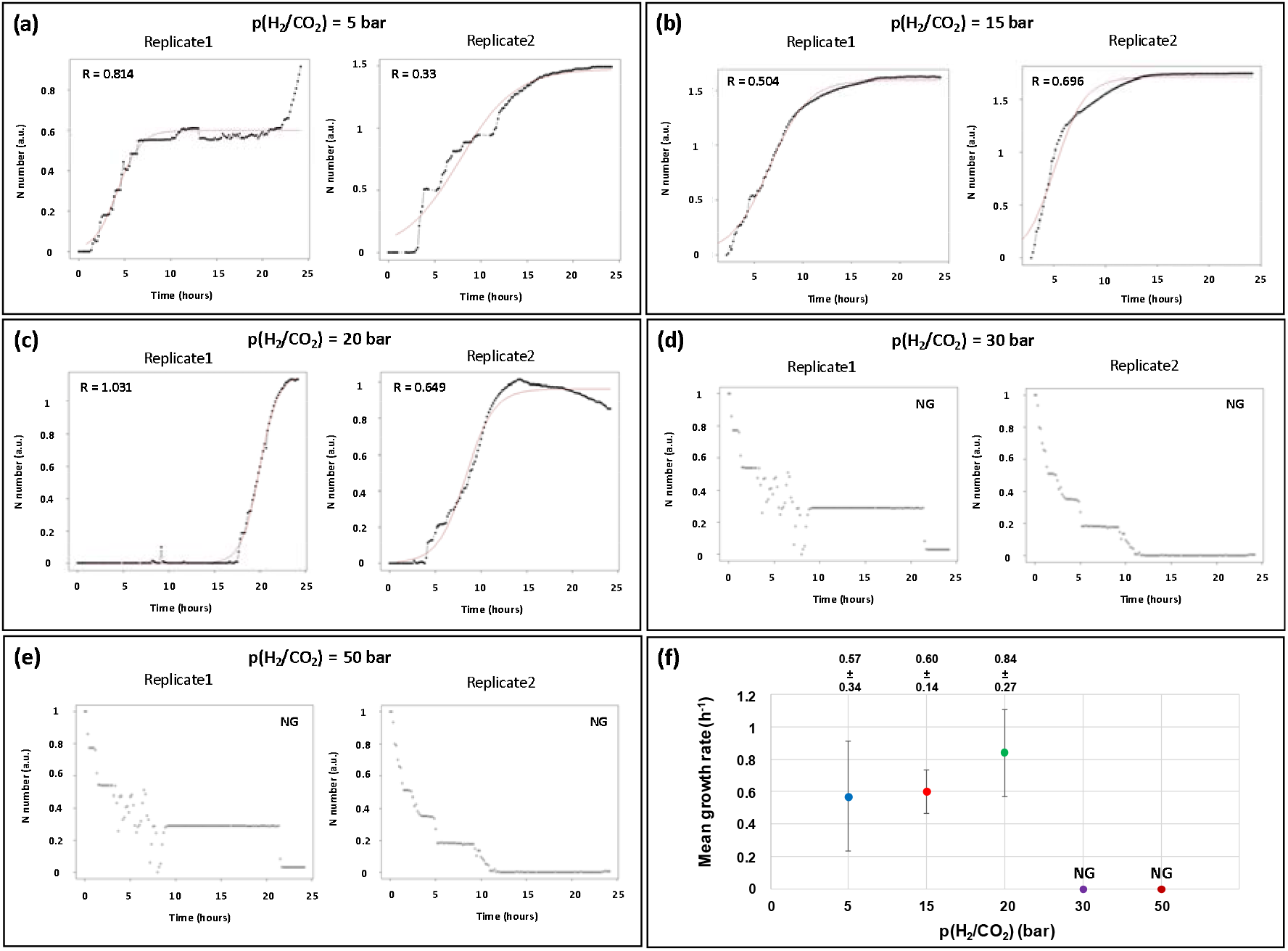
(a-e) Specific growth rates (R) calculated for each replicate across various p(H_2_/CO_2_) conditions (only the average growth rate is plotted). No growth (NG) was observed at p(H_2_/CO_2_) = 30 or 50 bar. (f) Mean growth rates of M. thermolithotrophicus under applied p(H_2_/CO_2_) (80/20 mol%) at a total pressure of 100 bar.

To better simulate conditions in deeper underground environments, stirring was removed, leading to a noticeable change in the strain behavior. The typical growth curves could no longer be measured via optical fibers, as a significant biofilm structure developed on the tube wall (Figure 5). This biofilm formation hindered the correlation of OD variation with cell density. Although planktonic cells were detected in the liquid phase, their concentration never exceeded 1·10^7^ cells.mL^-1^ (approximately an OD of 0.53). Consequently, estimating cell development under these conditions has become challenging. Notably, no biofilms were observed when the system was stirred, indicating that the shear forces generated during stirring prevented cell attachment and aggregation. Instead, biofilm formation was likely driven by local gas concentrations due to the concentration gradients created by slow gas diffusion and potential toxicity effects. Biofilm formation was evident across all the tested partial pressures, with the distance from the biofilm to the gas⍰liquid interface ranging from 1 ± 0.5 mm to 17 ± 2 mm (p(H_2_/CO_2_) from 5 to 30 bar) and the thickness varying between 0.1 ± 0 mm and 2.3 ± 0.5 mm.

**Figure 5.**
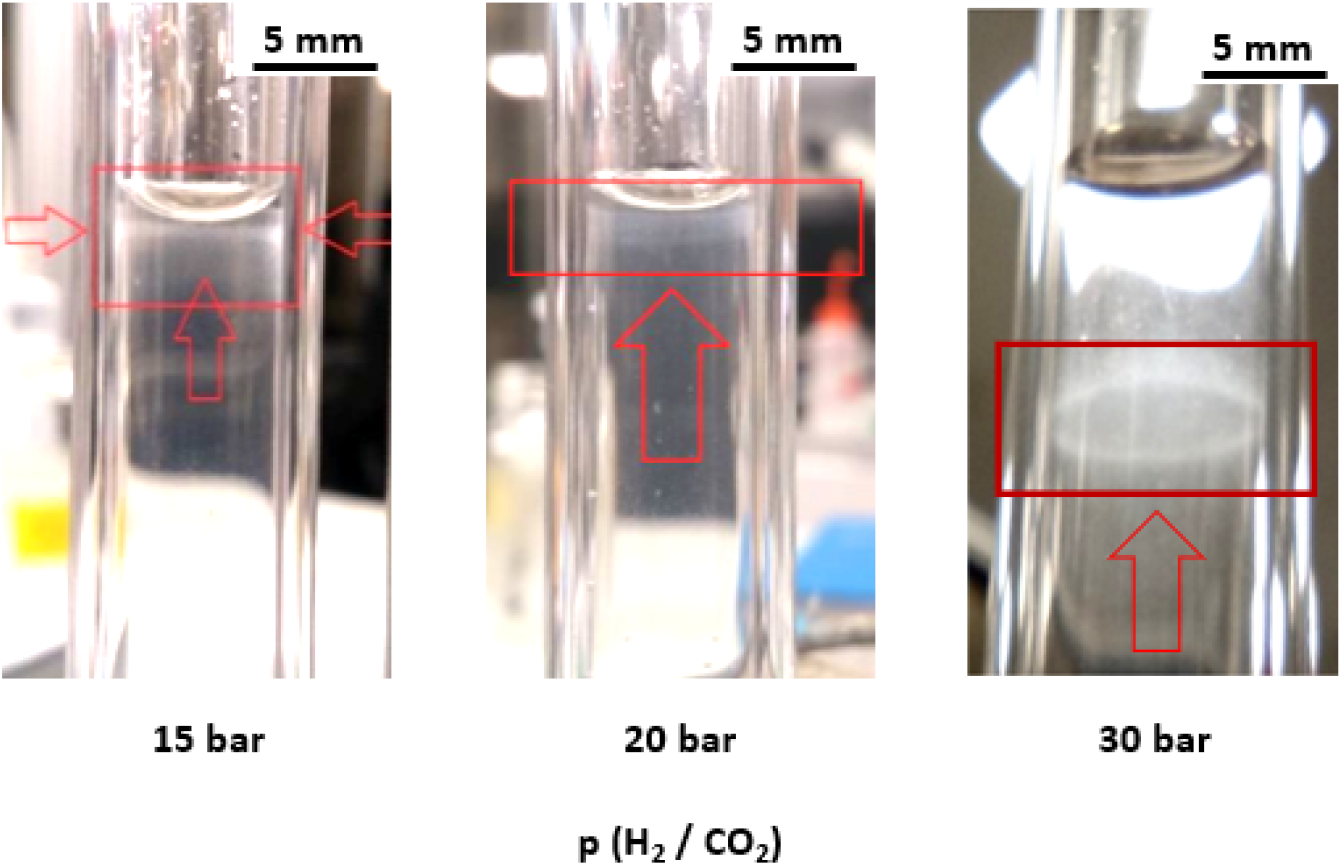
Images of biofilms of M. thermolithotrophicus inside the HPTR at a total pressure of 100 bar and various p(H_2_/CO_2_) ratios (80/20 mol%). At a pressure of 30 bar for p(H_2_/CO_2_), the development of the biofilm can be distinctly observed in a halo pattern on the wall of the high-pressure reactor (HPTR).

#### 2.12. Methane Production

Methane production was measured under each p(H_2_/CO_2_) condition, both with and without stirring (Figure 6). A significant observation was that methane production varied by at least an order of magnitude, being 8 to almost 1000 times higher depending on the applied p(H_2_/CO_2_) when stirring was absent. Under stirred conditions, methane production was limited to p(H_2_/CO_2_) = 20 bar, with the highest production reaching 8.6·10^−7^ ± 5.10^−7^ mol at p(H_2_/CO_2_) = 15 bar, which was correlated with the peak growth rate observed. Beyond p(H_2_/CO_2_) ≥ 30 bar, neither methane production nor cell growth was detectable.

**Figure 6.**
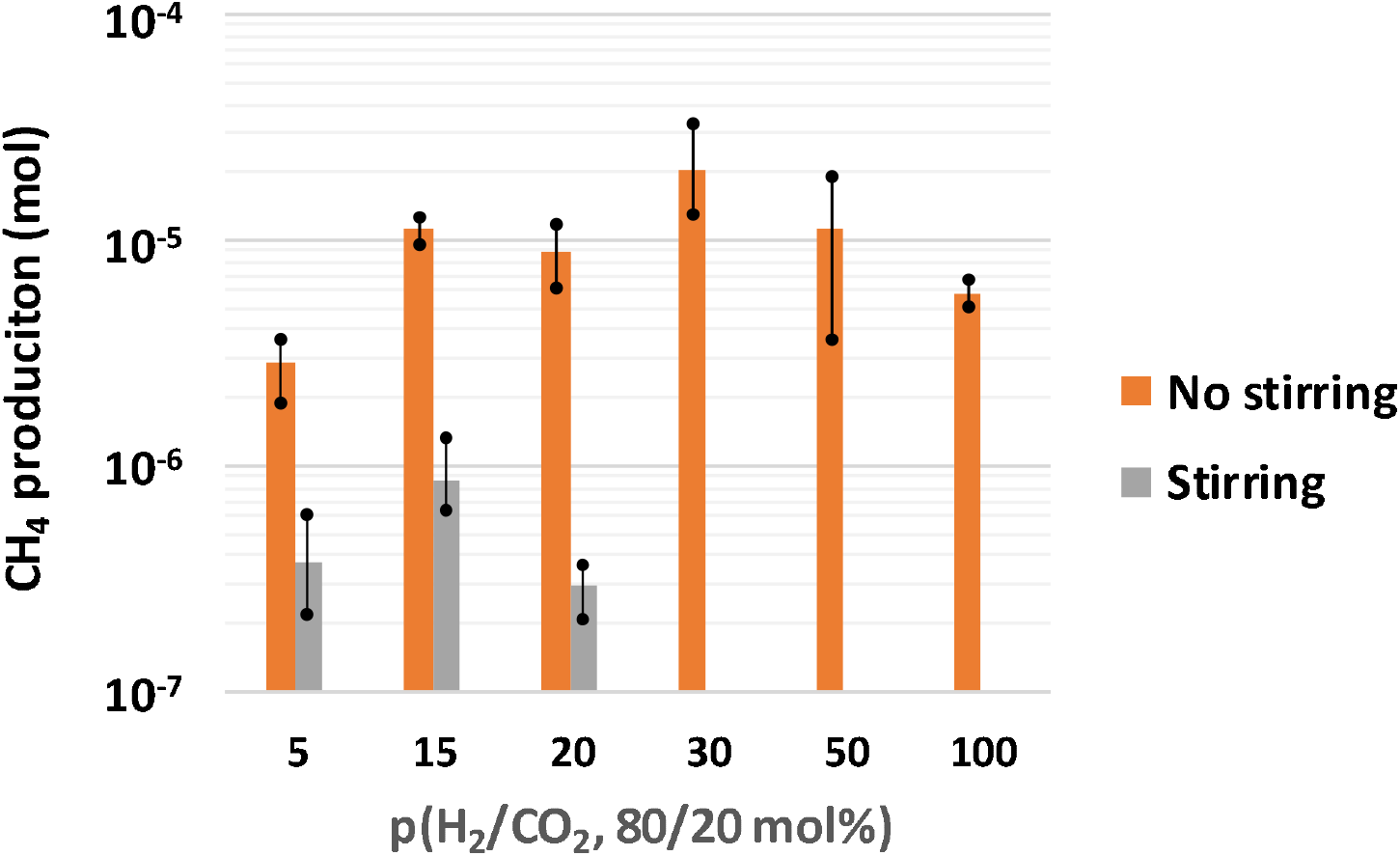
Methane production by M. thermolithotrophicus over 24 h at 65°C, a total pressure of 100 bar, and various p(H_2_/CO_2_) (80/20 mol%) with and without stirring (log scale).

In contrast, methane production occurred across all the tested conditions when stirring was removed, reaching 2.07·10^−5^ ± 5.60·10^−6^ mol at p(H_2_/CO_2_) = 30 bar before decreasing at higher pressures. This increase in production may be linked to biofilm formation, which tends to increase metabolic activity.

The presence of biofilms has been associated with increased methane productivity in biomethanation studies^49–51^. Notably, biofilms of methanogenic communities are predominantly composed of hydrogenotrophic species, which exhibit increased methane production when only biofilm cells are present in the environment^49^. The authors reported that hydrogenotrophic species could constitute up to 37.9% of the community, highlighting their prominence^49^. Furthermore, Maegaard and colleagues^50^ demonstrated that H^2^ flux within biofilms surpassed equivalent liquid conditions in control tests, supporting the notion that biofilm structures can lead to increased methane production when a biofilm forms, which was confirmed in this study. The authors also reported that the production of methane varied from 7.2·10^−2^ to 1.68·10^−1^ L_CH4_.L_medium_^-1^.day^-1^ in their bioreactor, which contained a microbial community arranged into a biofilm, whereas the production of methane from 2·10^−2^ to 5·10^−2^ L_CH4_.L_medium_^-1^.day^-1^ was observed in the control. Consistent with these findings, our results indicate significantly greater CH_4_ production when a biofilm formed, with levels reaching 3.9·10^−1^ ± 9·10^−2^ L_CH4_.L_medium_^-1^.day^-1^ for p(H_2_/CO_2_) = 30 bar (Figure 5 and Table 1). Moreover, biofilm formation appears to enhance the strain’s tolerance to elevated H_2_ and CO_2_ concentrations, as methane was produced at these concentrations in the presence of a biofilm, whereas no production occurred at p(H_2_/CO_2_) above 20 bar (*i.e.*, 30 and 50 bar) under stirring conditions when biofilms did not form (Figure 6).

**Table 1.** Daily methane production per liter of medium by M. thermolithotrophicus at 65°C and a total pressure of 100 bar over 24 h, comparing conditions with and without stirring (biofilm formation).

| p(H<sub>2</sub>/CO<sub>2</sub>)<br>(80/20 mol%) | Produced methane – no stirring<br>(L<sub>CH<sub>4</sub></sub>·L<sub>medium</sub><sup>-1</sup>) in 24 hours | Produced methane – stirring<br>(L<sub>CH<sub>4</sub></sub>·L<sub>medium</sub><sup>-1</sup>) in 24 hours |
| --- | --- | --- |
| 5 | 4.6·10<sup>-2</sup> | 5.92·10<sup>-3</sup> |
| 15 | 1.78·10<sup>-1</sup> | 1.376·10<sup>-2</sup> |
| 20 | 1.42·10<sup>-1</sup> | 4.64·10<sup>-3</sup> |
| 30 | 3.31·10<sup>-1</sup> | 0 |
| 50 | 1.79·10<sup>-1</sup> | 0 |
| 100 | 9.3·10<sup>-2</sup> | 0 |

The differences in methane production can be attributed to the equilibrium maintained under stirred conditions, where H_2_ and CO_2_ concentrations are uniformly distributed in the liquid phase, potentially exerting toxic effects. In contrast, without stirring, the toxic effects develop more slowly, allowing the strain to expand in time for growth and metabolic activity. Notably, the most favorable pressure for metabolic activity increases from 15 bar with stirring to 30 bar without stirring, which aligns with the hypothesis regarding diffusion time. While biofilm formation prevented accurate measurements of the cell concentration and individual methane production per cell under nonstirred conditions, such data could be calculated during stirring. Therefore, under stirring conditions, the methane production per cell for p(H_2_/CO_2_) = 5, 15 and 20 bar was estimated to be approximately 1.1·10^−5^ nmol CH_4_.cell^-1^, 1.5·10^−5^ nmol CH_4_.cell^-1^ and 4.3·10^−5^ nmol CH_4_.cell^-1^, respectively This finding indicates that individual cellular production remains within a similar order of magnitude across all conditions, with a slight increase observed at p(H_2_/CO_2_) = 20 bar. This pattern suggests that as the pressure of H_2_/CO_2_ increases, the cells may begin to experience stress conditions, favoring metabolic activity over cellular proliferation. Importantly, even with improved access to and concentration of substrates, these factors do not enhance the overall production capability of the cells.

To further examine the behavior of methane production without stirring under optimal metabolic conditions (*i.e.*, p(H_2_/CO_2_) = 30 bar), production was monitored over a 24-hour period (Figure 7). Methane production exhibited linear and stable accumulation over time (R^2^= 0.98). The continued increase in production (accumulation of methane in the reactor) beyond the 24-hour mark can be attributed to the constant connection of the reactor to a high-pressure ISCO pump (260 mL), which was kept filled with the gas mixture (p(H_2_/CO_2_) (80/20 molar) = 30 bar, supplemented with nitrogen to achieve a total pressure of 100 bar). Consequently, the H_2_ and CO_2_ concentrations remained stable throughout the incubation period (approximately 260 mL of gas was added to the 1.5 mL of culture medium inside the HPTRs). This approach was designed to simulate underground hydrogen storage (UHS) and underground methane reactor (UMR) scenarios for storing millions of cubic meters of gas. The measurements obtained here are encouraging for stimulating methanogenic activity under a continuous supply of substrates (*i.e.*, UMR). Hypothetically, lithoautotrophic methanogenic archaea should be capable of producing indefinitely with a constant influx of substrates in the natural environment if we abstract from all other limiting nutrients.

**Figure 7.**
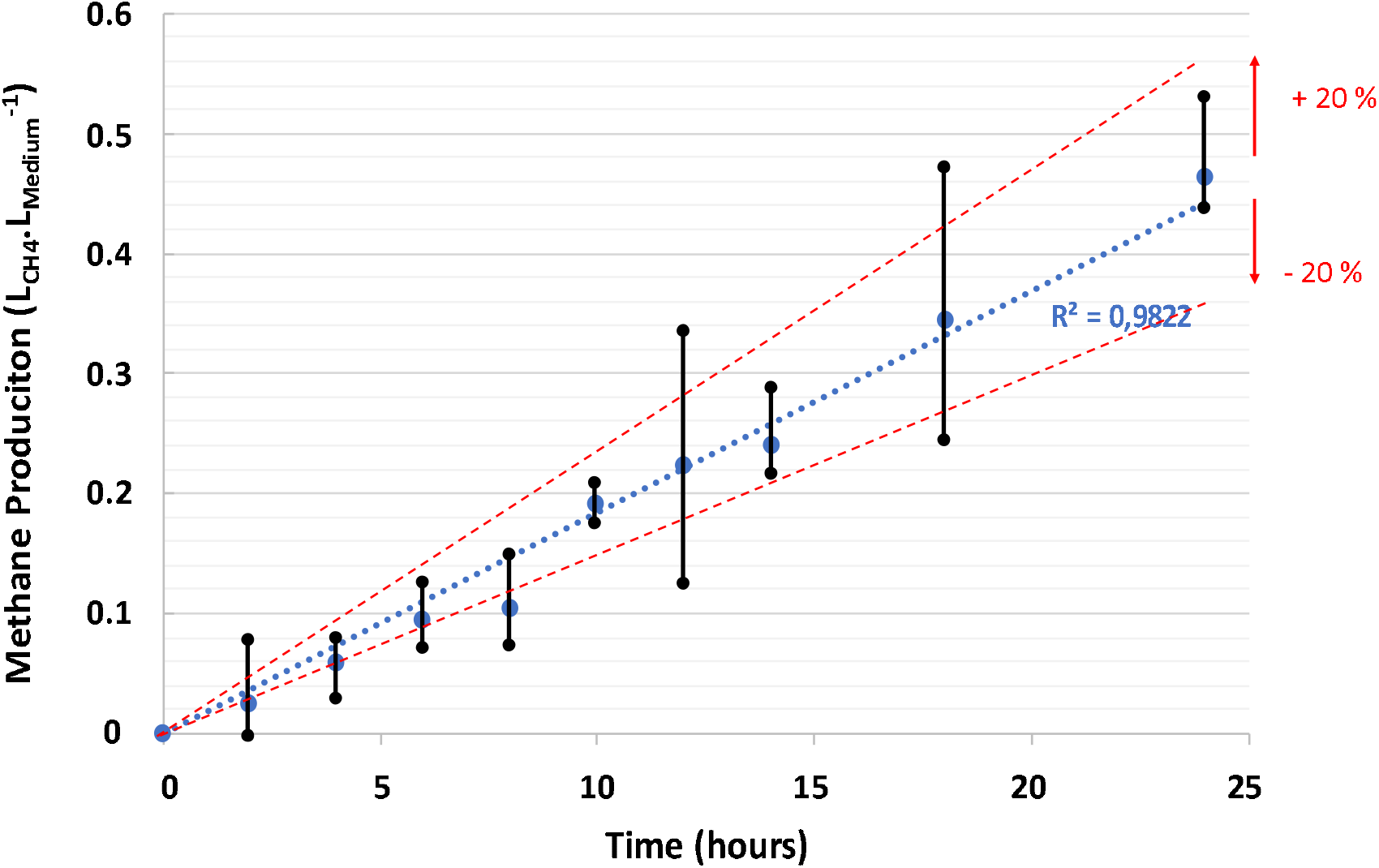
Methane production by M. thermolithotrophicus over 24 h at 65°C, a total pressure of 100 bar, and p(H_2_/CO_2_) (80/20 mol%) = 30 bar without stirring.

Under these optimized conditions (at 65°C, a total pressure of 100 bar and p(H_2_/CO_2_) = 30 bar), the methane production rate for *M. thermolithotrophicus* was found to be 1.85·10^−2^ L_CH4_.L_medium_^-1^. hour^-1^, equating to 4.4·10^−1^ ± 9·10^−2^ L_CH4_.L_medium_^-1^.day^-1^.

The kinetics of production also allowed the detection of biofilm formation, which began to appear on the tube walls between 2 and 4 hours of incubation. This formation period is notably rapid compared with similar observations reported under controlled laboratory conditions with model strains^52,53^. For example, compared with other model strains, *Pseudomonas aeruginosa* biofilm formation is often detected between 12 and 24 hours^52^ but varies between 24 and 72 hours for certain *Staphylococcus* species^53^. These findings suggest that *M. thermolithotrophicus* has an exceptional ability to develop biofilm structures under specific conditions.

### 2.2. Gas Solubility and Diffusion

Table 2 presents the concentrations of H_2_ and CO_2_ in the liquid phase, reflecting the experimental conditions. Although CO_2_ is present in a lower proportion than H_2_ in the gas mixture (80% H_2_ to 20% CO_2_), our calculations indicate that CO_2_ concentrations in the liquid phase were higher, with a H_2_/CO_2_ molar ratio largely in favor of CO_2_.

**Table 2.**
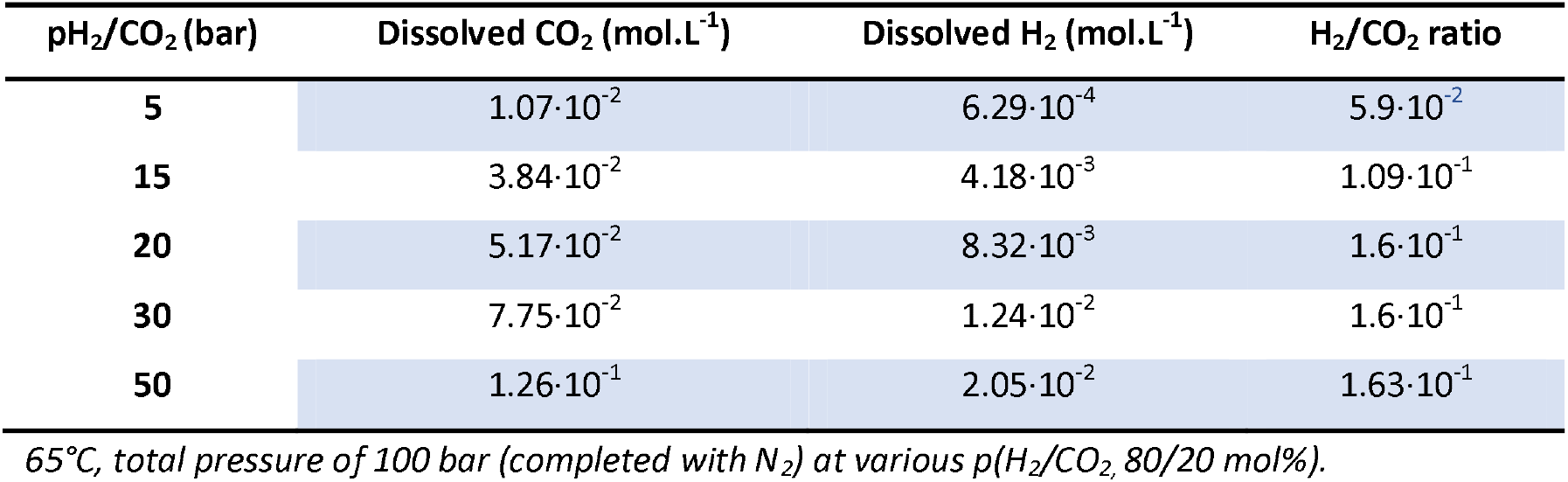
Dissolved CO_2_ and H_2_ in the growth medium under the explored experimental conditions:

As anticipated, both gases exhibited increased solubility with increasing partial pressures in the culture medium. However, CO_2_ demonstrated higher solubility than H_2_. At the maximum p(H_2_/CO_2_) of 50 bar, the concentration of CO_2_ in the liquid phase reached 1.26·10^−1^ mol.L^-1^, whereas H_2_ solubility was much lower at 2.05·10^−2^ mol.L^-1^. Owing to its small molecular size, H_2_ is more volatile and challenging to dissolve. The influence of ions in saline solutions can complicate dissolution, as indicated in our calculation, which corroborates findings from previous studies^54^. The H_2_/CO_2_ ratio in the liquid phase increased from 5.9·10^−2^ at 5 bar to 1.63·10^−1^ at 50 bar, displaying a rapid increase with increasing p(H_2_/CO_2_) followed by stabilization above 20 bar. This suggests that H_2_ solubility increases more significantly than CO_2_ solubility with increasing partial pressure. Notably, the observed tolerance of *M. thermolithotrophicus* to elevated p(H_2_/CO_2_), inferred from its growth data and methane production rates, peaked between 20 and 30 bar, which coincided with the stabilization of the H_2_/CO_2_ ratio. Beyond p(H_2_/CO_2_) = 30 bar, methane production decreased, indicating that the strain’s tolerance to H_2_ and CO_2_ may be exceeded even when biofilm formation occurs.

Furthermore, the diffusion of gases in the medium over time is crucial and reveals distinct kinetic behaviors for each gas under purely diffusive scenarios (*i.e.*, without stirring and neglecting convection). As shown in Figure 8, H_2_ diffused more rapidly than CO_2_ across all the conditions examined. Even after 13 hours of incubation, the system had not achieved full equilibrium, indicating that both gases remained more concentrated near the gas/liquid interface. Our simulations indicate that the H_2_ concentration at the biofilm level after one hour of incubation (at a depth of 17 ± 2 mm for p(H_2_/CO_2_) = 30 bar) was 5·10^−4^ mol.L^-1^, whereas the CO_2_ concentration was significantly greater at 1.75·10^−2^ mol.L^-1^. This localized concentration of H_2_ and/or CO_2_ (above p(H_2_/CO_2_) = 20 bar). In contrast, under nonstirred conditions, biofilm formation appears to provide a protective mechanism, enabling the strain to withstand the harmful effects of H_2_ and increasing methane production yields compared with those lower p(H_2_/CO_2_) conditions (Figure 6). In a stirred system, equilibrium conditions render the concentrations of H_2_ and CO_2_ uniform throughout the reactor, whereas nonstirred conditions facilitate varying concentrations and diffusion kinetics on the basis of depth from the interface.

**Figure 8.**
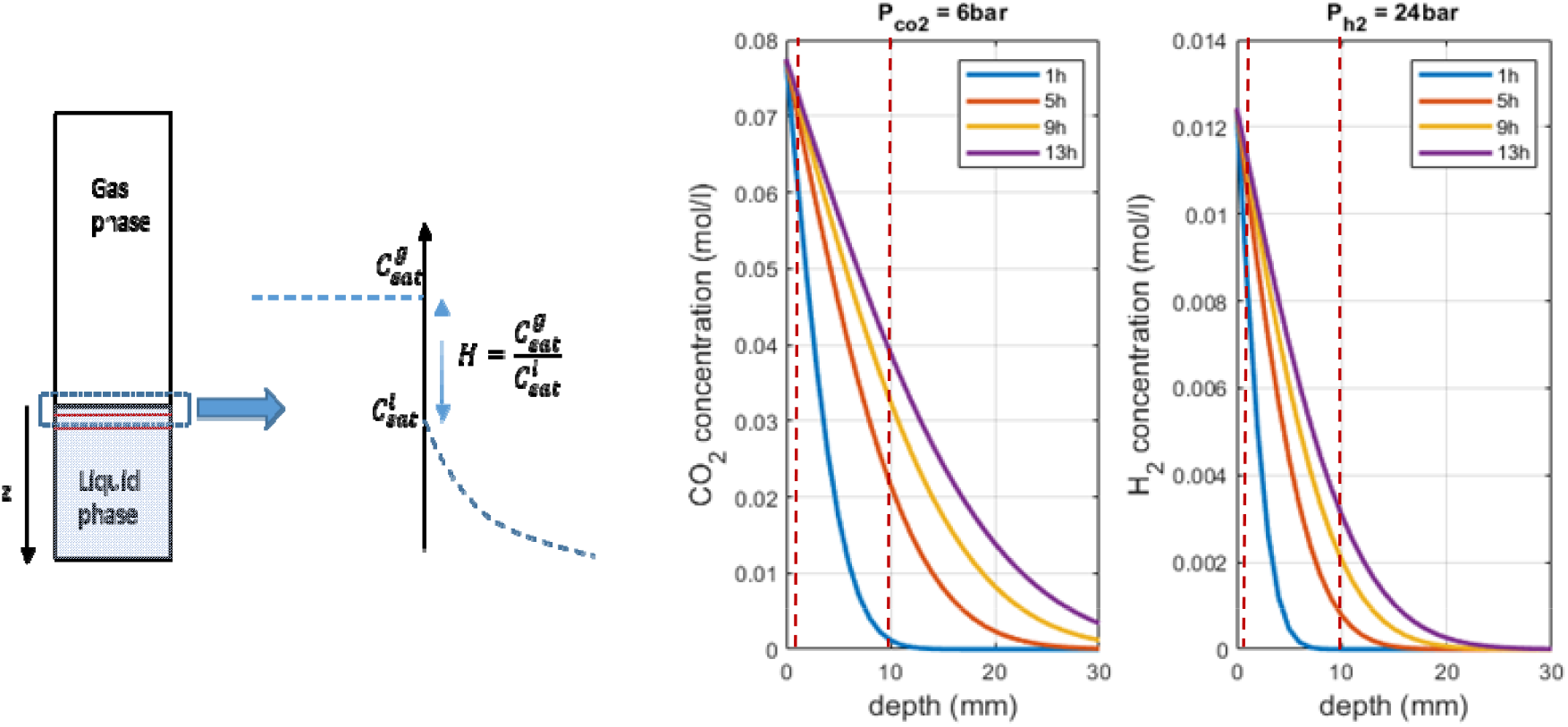
CO_2_ and H_2_ concentrations in the medium as a function of the distance (depth) from the gas–liquid interface over time for T = 65°C and total pressures of 100 bar (supplemented with nitrogen) and p(H_2_/CO_2_) (80/20 mol%) = 30 bar.

Our calculations confirm that CO_2_ demonstrates higher overall solubility, although H_2_ solubility increases more rapidly. CO_2_ can affect cell metabolism through environmental acidification, leading to pH levels that may hinder cellular development^41^. However, pH measurements before and after each experiment, as presented in ESI-2, revealed a slight decrease at high partial pressures, stabilizing at approximately pH ≈ 5.7–5.9 (down from 6.5 prior to gas injection) due to the presence of HEPES buffer. These pH values remain within a favorable range for metabolic activity and cannot account for the observed growth variations, as previously reported^41^. Additionally, high concentrations of CO_2_ can be detrimental to cells. For example, supercritical CO_2_ is known to have sterilizing properties, disrupting membrane integrity and leading to cell death^55^. Nevertheless, in our experimental setup, CO_2_ was diluted, among other gases, which mitigated its toxic effect. Previous studies evaluating *M. thermolithotrophicus* CO_2_ tolerance revealed detrimental impacts on metabolic activity at elevated pressures, particularly in research conducted by Dupraz and colleagues^41^. They observed reduced metabolic function when CO_2_ concentrations increased, with experiments conducted at pressures up to 10 bar and varying H_2_/CO_2_ ratios, all using the same culture medium as our experiments, *i.e.*, AGW, with NaOH as a buffer. The calculated values of dissolved gases under their experimental conditions (see ESI-3) were 2.13·10^−4^ < [CO_2_] (mol.L^-1^) < 9.69·10^−2^ and 6.02·10^−4^ < [H_2_] (mol.L^-1^) < 5.45·10^−3^, with p(H_2_/CO_2_) ranging from 1.1 to 10 bar. Under our conditions, metabolic activity begins to decrease for p(H_2_/CO_2_) > 30 bar, at which point the CO_2_ concentration at equilibrium in the medium is 7.75·10^−2^ mol.L^-1^, which is quite close to the level reported by Dupraz and collaborators for p(H_2_/CO_2_) = 10 bar.

Another potential contributor to the observed effects may be the toxicity of H_2_ itself. Prior investigations have shown that high concentrations of H_2_ can inhibit cell growth in nonmethanogenic strains^56,57^, whereas biofilm formation in methanogens seems to improve when H_2_ exposure in the liquid phase is minimized^49^. Jensen and colleagues^49^ reported that reducing the H_2_ retention time can significantly increase biofilm activity and methane production in methanogenic communities, resulting in up to 12.5 times greater effects at the lowest retention time than at the highest retention time. Similarly, research conducted in anaerobic digesters at 35°C and ambient pressure (up to 1.55 bar) has demonstrated that H_2_ pressure beyond 0.8–0.9 bar negatively affects cumulative methane production, indicating a threshold above which methane-generating processes can be severely impaired. H_2_ was consistently consumed during the reactions; however, at p(H_2_) = 1.557 bar, methanogenesis was significantly hindered, and hydrolysis/acidogenesis ceased. These results of the deleterious effects of high H_2_ partial pressure were notably supported by the fact that CO_2_ was absent under most of the tested conditions and that when CO_2_ was added to the system, metabolism was disrupted again. Indeed, for the same total pressure, *i.e.*, p(H_2_) = 1.55 bar, but under a H_2_/CO_2_ atmosphere, the methanogenic activity was greater than that under only H_2_ conditions^58^. Despite the fact that all of these findings were made for ambient pressure conditions, the results that we obtained under much higher-pressure conditions seem to agree with the general trend observed in these studies: our system did present greater methane production when the strain exhibited biofilm behavior, as well as a decrease in metabolic activity and growth above a certain threshold. The potential negative impact of dihydrogen could then be assessed for methanogens in the frame of geological storage of this molecule^13^, even if its injection in underground formations could be favorable to other kinds of microorganisms, which could present greater tolerance. The fact that more H_2_ is needed than CO_2_ for methanogenesis leads to a faster decrease in its concentration in laboratory-controlled investigations; however, under high partial pressure, for example, in the case of UHS and UMR, it can be hypothesized that consumption is not fast enough to counteract the deleterious impact. The cells will then be quickly impacted; their metabolic activity will be reduced, leading to a longer accumulation of H_2_ in the environment, leading to deleterious impacts on the cells and so on.

One last hypothesis could be that both gases may reach toxic concentrations under elevated pressure conditions (*i.e.*, above 30 bar), thereby diminishing growth and metabolic performance. A hypothetical synergistic toxicity could emerge from simultaneous exposure to both H_2_ and CO_2_, even if neither is present at their maximum tolerable levels. This dual stress could demand excess energy for cellular maintenance and metabolism, consequently lowering the strain’s resilience as H_2_ and CO_2_ pressures increase.

Finally, biofilm formation presents significant potential in the context of geological CO_2_ storage, particularly for UMR, primarily through its effects on safety and production yields. The presence of biofilms within rock pores can increase sequestration safety by altering the permeability of the reservoir and sealing fractures, which could facilitate CO_2_ leakage, a phenomenon that has been supported by the literature^59–61^. Additionally, the presence of biofilms is correlated with increased biovalorization and energy production. However, concerns regarding decreased injectivity due to bioclogging during fluid storage or withdrawal are commonly raised. In current UGSs, bioclogging near injection and production wells is deemed unlikely for two reasons: (i) direct contact with gases at such high pressures is a powerful biocide and (ii) massive gas injection over years, or even decades, has most certainly dried out the rock, making microbial difficult to settle. However, biofilm formation may still occur in surrounding environments, promoting beneficial processes that valorize stored CO_2_. Environmental biofilms may harbor fermenters capable of producing H_2_ and enhancing the methanogenesis reaction through metabolic cooperation.

## 3. Conclusion

The primary objective of this research was to determine how the partial pressures of H_2_ and CO_2_ influence the growth and yield of methanogenesis. To achieve this goal, *Methanothermococcus thermolithotrophicus* was cultivated in innovative sapphire-based high-pressure transparent reactors (HPTRs) that allow continuous monitoring of growth at a pressure of 100 bar, which is representative of underground gas storage (UGS) conditions where CO_2_, H_2_, or both can be injected. Notably, the absence of stirring in these reactors, which more accurately simulate natural underground conditions, resulted in the formation of biofilms that significantly increased methane production across the various investigated p(H_2_/CO_2_, 80/20 mol%) partial pressures. These findings underscore the critical role of biofilm dynamics in methanogenic processes. Furthermore, the complex interactions between H_2_ and CO_2_ at elevated pressures raise important considerations regarding the potential toxic effects and metabolic stress on microbial populations, challenging existing assumptions about gas interactions in high-pressure environments. The obtained results provide valuable insights into optimizing methanogenic pathways in the context of carbon capture and storage, as well as the sustainable management of underground gas reservoirs.

## Supporting information

ESI-1

ESI-2

ESI-3

## Declarations

### Ethics approval and consent to participate

Not Applicable

### Consent for publication

Not Applicable

### Availability of data and materials

The datasets generated during and/or analysed during the current study are available in the Electronic Supporting Information (ESI-1 – 3).

## Authors’ contributions

E.V., A.C., A.R-P and S.M. designed the work; E.V., A.C., M.J., O.N. participated in the acquisition of the data; E.V., A.C., A.E., A.R-P and S.M. interpreted the data, E.V. drafted the main text, E.V., A.C., O.N., A.E., A.R-P and S.M. made the figures, A.C., A.E., A.R-P and S.M. finalized and revised the work.

## Acknowledgments

The authors would like to thank Marion Guignard (Université de Pau et des Pays de l’Adour, CNRS, IPREM) for providing trace element stock solutions for Archean cultures. Fabien Palencia (ICMCB, CNRS) is also acknowledged for the design and 3D printing fabrication of the optical fiber supports.

