## Supplementary material for "Real time monitoring of hydrogenotrophic methanogenesis under deep saline aquifers conditions": ESI-2

**Evolution of the pH of the AGW medium as a function of the H_2_/CO_2_ partial pressure**

The pH of the AGW growth medium was measured with a pHmeter (HANNA Instruments, pH HI 2210 pH meter) once the system is decompressed.

Although the medium was supplemented with a buffer (HEPES, 120 mM), the pH is decreasing down to 5.6 when the pressure is reaching p(H_2_/CO_2_) = 100 bar, while it is initially set around 6.5.

However, the buffer is efficient as the pH can be stabilized at this value and remain between 5.5 and 6 for all of the measured conditions (Figure ESI2-1). Despite these slightly acidic conditions, it remains in the viable zone for our two selected strains, suggesting that they should be able to proliferate and have an active metabolism.

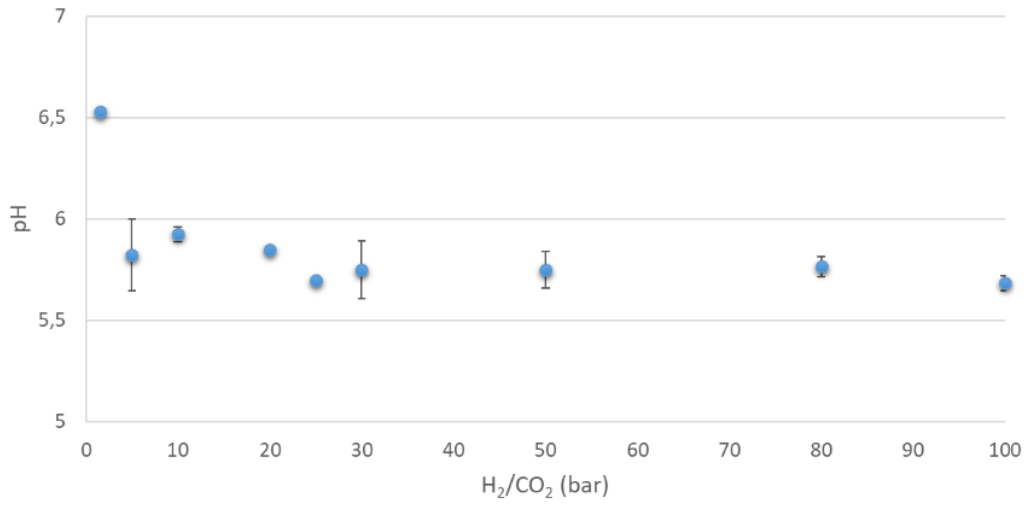

**Figure ESI2-1.** Ex situ measurement of pH value of AGW media after pressurising it with H_2_ /CO_2_ (4:1).
