## Supplementary material for "Real time monitoring of hydrogenotrophic methanogenesis under deep saline aquifers conditions": ESI-3

**Calculated values of dissolved gases under the experimental conditions of Dupraz *et al.***

**Table 2.** Calculated dissolved H_2_ and CO_2_ in the experimental conditions of Dupraz et al.: Dupraz, S.; Fabbri, A.; Joulian, C.; Dictor, M.-C.; Battaglia-Brunet, F.; Menez, B. Impact of CO2 Concentration on Autotrophic Metabolisms and Carbon Fate in Saline Aquifers–A Case Study. Geochim. Cosmochim. Acta 2013, 119, 61–76.

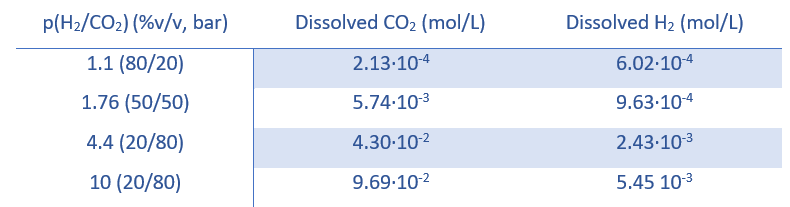
